# Deciphering anti-infectious compounds from Peruvian medicinal *Cordoncillos* extract library through multiplexed assays and chemical profiling

**DOI:** 10.1101/2022.11.15.516654

**Authors:** Pedro G. Vásquez-Ocmín, Sandrine Cojean, Vincent Roumy, Guillaume Marti, Sébastien Pomel, Alice Gadea, Karine Leblanc, Indira Dennemont, Liliana Ruiz-Vásquez, Hivelli Ricopa Cotrina, Wilfredo Ruiz Mesia, Stéphane Bertani, Lastenia Ruiz Mesia, Alexandre Maciuk

## Abstract

High prevalence of parasitic or bacterial infectious diseases in some world areas is due to multiple reasons, including a lack of an appropriate health policy, challenging logistics and poverty. The support to research and development of new medicines to fight infectious diseases is one of the sustainable development goals promoted by World Health Organization (WHO). In this sense, the traditional medicinal knowledge substantiated by ethnopharmacology is a valuable starting point for drug discovery. This work aims at the scientific validation of the traditional use of *Piper* species (“*Cordoncillos”*) as firsthand anti-infectious medicines. For this purpose, we adapted a computational statistical model to correlate the LCMS chemical profiles of 54 extracts from 19 *Piper* species to their corresponding anti-infectious assay results based on 37 microbial or parasites strains. We mainly identified two groups of bioactive compounds (called features as they are considered at the analytical level and are not formally isolated). Group 1 is composed of 11 features being highly correlated to an inhibiting activity on 21 bacteria (principally Gram-positive strains), one fungus (*C. albicans*), and one parasite (*Trypanosoma brucei gambiense*). The group 2 is composed of 9 features having a clear selectivity on *Leishmania* (all strains, both axenic and intramacrophagic). Bioactive features in group 1 were identified principally in the extracts of *Piper strigosum* and *P. xanthostachyum*. In group 2, bioactive features were distributed in the extracts of 14 *Piper* species. This multiplexed approach provided a broad picture of the metabolome as well as a map of compounds putatively associated to bioactivity. To our knowledge, the implementation of this type of metabolomics tools aimed at identifying bioactive compounds has not been used so far.

## INTRODUCTION

In Amazonia, tropical diseases have a high prevalence due to multiple reasons, including a lack of an appropriate health policy, a climate conducive to diseases, challenging logistics and poverty. Neglected tropical diseases (NTDs) are a group of 20 conditions (caused by viruses, protozoa, helminths and bacteria) prioritized by the World Health Organization (WHO). The NTDs are responsible for approximately 200,000 deaths and the loss of 19 million disability-adjusted life years (DALYs) annually. In 2020, new infections by *Plasmodium* spp. and *Leishmania* spp. worldwide were estimated at 241 and 1 million, respectively. Even if *Plasmodium* is not strictly classified as a NTD, it caused around 627,000 deaths in 2020, with two-thirds of these deaths (470,000) being due to treatment disruptions during the COVID-19 pandemic (WHO, 2021). *Leishmania* spp. and *Trypanosoma* spp. are protozoan parasites responsible of diseases labeled as NTDs *stricto sensu*. The ambitious 10-year WHO plan to defeat NTDs is based on three pillars: control, elimination and eradication (WHO, 2022a). The support of research and development of new medicines to fight infectious diseases is one of the sustainable development goals promoted by WHO (WHO, 2022b). Drug resistance is a widespread concern in medical care, and the increase of drug-resistant infections is faster than the pace of the development of new drugs approved for use in humans. Therefore, every input into the search for new antimicrobial agents is welcome (Yan et al., 2021). Ethnopharmacology is the interdisciplinary study of the knowledge or practices of traditional cultures related to plants, animals, or mineral used for therapeutic purposes (SFE, 2022). Such knowledge can be valued as the starting point for drug discovery, especially in the case of infectious diseases which are prevalent among such populations.

Piperaceae is a pantropical family composed of eight genera. Two genera are the most representative in this family: *Piper* and *Peperomia* (Vásquez-Ocmín et al., 2017; Salehi et al., 2019). Among these, *Piper* is the most diverse and representative genus, encompassing *ca*. 2600 species (Trujillo et al., 2022). Besides pungent compounds, many *Piper* species produce essential oils and are hence highly aromatic, explaining their use for cooking and medicinal purposes (Ruiz-Vásquez et al., 2022). In Peru, *Piper* spp. (called “Cordoncillos”) have been used for a very long time in traditional medicine as a “first-hand treatment”, especially in the villages far away from major cities and medical care. The necessary scientific validation of traditional uses implies the isolation of the main bioactive compounds by successive fractionations using chromatographic techniques and biological activity testing (bio-guided isolation). Such an approach has evidenced interesting activities for *Piper* spp. as antiinflammatory, antiparasitic, antibacterial, etc. (Mgbeahuruike et al., 2017; Durant-Archibold et al., 2018). Even if bio-guided fractionation is still used with some success in the natural product chemistry field, current trends involve streamlining the cost, effort, and time (Vásquez-Ocmín et al., 2022a). Metabolomics is a holistic approach allowing rapid detection and putative identification (*i.e*. annotation) of numerous metabolites, along with data mining on multiple datasets. Metabolomics aims to comprehensively map all biochemical reactions in a given system and has become a key to deciphering their biological roles, hence becoming mainstream in natural product chemistry and drug discovery. This approach can be divided into targeted and untargeted analyses (Alarcon-Barrera et al., 2022). Two spectral techniques are regularly employed for metabolomics analysis: mass spectrometry (MS) and nuclear magnetic resonance (NMR) spectroscopy, both being assisted by bioinformatics and statistical analysis. Our team has solid expertise in implementing straightforward and effective workflows to decipher anti-infectious compounds using untargeted metabolomics (Vásquez-Ocmín et al., 2021a).

This work aims at the scientific validation of the use of *Piper* species as firsthand anti-infectious medicines. We adapted a statistical model to correlate the LCMS chemical profiles of 54 extracts from 19 *Piper* species to their corresponding anti-infectious assay results based on 37 microbial or parasites strains. This multiplexed approach led to the annotation of compounds bearing these activities.

## MATERIALS AND METHODS

### Ethnopharmacology and Plant Material

Based on the encouraging previous results of our research group on the anti-infectious activities of *Piper* species (Vásquez-Ocmín et al., 2021a), ethnopharmacological surveys were realized as part of the project “Compuestos bioactivos *in vitro* a partir de especies vegetales Amazónicas”. The surveys were undertaken between July and December 2020 in communities of three Amazonian regions of Peru: Cusco, Loreto and San Martin. People of these communities were interrogated about the main use of medicinal *Piper* species (“*Cordoncillos*”), including their use against malaria, “uta” (local name for leishmaniasis) and bacteria. According to ethnopharmacological studies, we collected 8 species in Cusco, 10 species in Loreto, and 1 species in San Martin.

Thus, different parts of these nineteen plants were collected, then identified and deposited in the Herbarium Amazonense (AMAZ), Iquitos, Peru. This project was realized in accordance with the guidelines pertaining to ethnopharmacological studies and edited by the Laboratorio de Investigacion de Productos Naturales Antiparasitarios de la Amazonia (LIPNAA) of the Universidad Nacional de la Amazonia Peruana (UNAP) (Resolucion Rectoral N° 1312-2020-UNAP).

### Preparation of plant extracts

One hundred grams of each air-dried and ground plant (leaves, leaves and stems, aerial parts) were soaked in 1 L of each of the following solvents successively: hexane, methylene chloride, methanol, ethanol/water (7:3 v/v), and aqueous. Extractions were made for 21 days for hexane, methylene chloride and ethanol/water, and for 15 days in methanol and aqueous extracts, with solvents being changed every 3 days. The extracts were then filtered through a paper filter and evaporated under reduced pressure below 40°C. Dry extracts were stored at −4°C until use. Extracts were solubilized in DMSO at a concentration of 10 mg/mL for *in vitro* bioassays and HPLC-MS analysis.

### Cell lines and microorganism culture

#### HUVEC cells

Human umbilical vein endothelial cells (HUVECs) were maintained in culture in RPMI 1640 medium (Invitrogen, Life Technologies) supplemented with 10 % heat-inactivated fetal bovine serum (Invitrogen Life Technologies) and 1 mM glutamine (Invitrogen Life Technologies) (Vásquez-Ocmín et al., 2018).

#### RAW264.7

The mouse monocyte/macrophage cell line RAW264.7 was maintained in culture in DMEM (Invitrogen, Life Technologies) supplemented with 10 % heat-inactivated fetal bovine serum (Vásquez-Ocmín et al., 2018).

#### Plasmodium

The *P. falciparum* chloroquine-sensitive strain 3D7 was obtained from the Malaria French National Reference Center (CNR Paludisme, Hôpital Bichat Claude Bernard, Paris) and was maintained in O^+^ human erythrocytes in RPMI 1640 medium (Invitrogen, Life Technologies) supplemented with 25 mM HEPES (Sigma), 25 mM NaHCO3 (Sigma) and 0.5 % Albumax II (Invitrogen, Life Technologies) at 37°C in a candle-jar method following the Trager and Jensen conditions (Trager and Jensen, 1976; Lambros and Vanderberg, 1979; Vásquez-Ocmín et al., 2018).

#### Leishmania

The *L. donovani* MHOM/ET/67/HU3, *L. amazonensis* MHOM/BR/73/M2269, and *L. braziliensis* MHOM/BR/75/M2903b strains were maintained routinely in *in vitro* culture. Passages in macrophages RAW 264.7 were carried out regularly to preserve the virulence of the strain, then they were recovered for culture maintenance in the promastigote form. For assays, parasites were maintained as promastigote forms in M-199 medium (Sigma) supplemented with 40 mM HEPES, 100 mM adenosine, 0.5 mg/L hemin and 10 % fetal bovine serum (FBS) at 25°C in a dark environment (Balaraman et al., 2015; Vásquez-Ocmín et al., 2018).

#### Trypanosomes

Trypomastigotes of *T. b. gambiense* (FéoITMAP/1893 strain) were grown in HMI9 medium constituted of prepacked Iscove’s modified Dulbecco’s medium (Thermo-Fisher, Les Ulis, France) supplemented with 36 mM NaHCO3, 1 mM hypoxanthine, 0.05 mM bathocuproine, 0.16 mM thymidine, 0.2 mM 2-mercapthoethanol, 1.5 mM L-cysteine, 10 % heat-inactivated foetal bovine serum, 100 IU penicillin and 100 μg.mL^-1^ streptomycin. Parasites were incubated in a Series 8000 direct-heat CO_2_ incubator (Thermo-Fisher, Les Ulis, France) at 37°C in a water-saturated atmosphere containing 5 % CO_2_ (Pomel et al., 2015; Vásquez-Ocmín et al., 2018).

#### Bacteria and yeast

Most microbial strains were diluted in BH medium (Brain Heart), MH medium (Mueller Hinton) for *Candida sp*. and *Mycobacterium sp*., or WW medium (Wilkins-West) for *Streptococcus sp*. and stored at −20°C. Then strains were subcultured at 20°C on RC (Ringer Cysteine) medium for 24 hours before tests (Bocquet et al., 2019).

### *In vitro* antiprotozoal activity

#### In vitro antiplasmodial activity on P. falciparum

Assays were realized with a suspension of erythrocytes at 1 % parasitemia containing more than 85 % ring stage obtained by repeated sorbitol treatment and incubated with the compounds at concentrations ranging between 0.49 and 100 μM or μg/mL, obtained by serial dilution, in duplicates. Two controls were used, parasites without drug and parasites with chloroquine at concentrations ranging between 0.49 and 1000 nM. Plates were incubated for 44 h at 37 °C in a candle jar (Vásquez-Ocmín et al., 2018, 2021a).

#### In vitro antileishmanial activity on L. donovani, L. amazonensis, and L. braziliensis axenic amastigotes

A suspension of promastigotes in growth plateau-phase was incubated at 37°C in 5 % CO_2_ for 3 days to obtain the amastigote form in promastigote medium supplemented with 2 mM CaCl_2_, 2 mM MgCl_2_, and a pH adjusted to 5.5. The axenic amastigote suspension containing 1.106 parasites/mL was incubated for 72h with the compounds at 37°C in 5 % CO_2_ in the dark. Tested compounds or extracts were obtained by serial dilution and ranged between 0.49 and 100 μM or μg/mL. There were two controls: parasites without drug and parasites treated with miltefosine at the same concentrations as the compounds tested (Vásquez-Ocmín et al., 2018, 2021a).

#### In vitro antileishmanial activity on L. donovani, L. amazonenesis and L. braziliensis intramacrophage amastigote form

Macrophages were seeded into a 96 well microtitration plate at a density of 100,000 cells/well in 100 μL and incubated in a 5 % CO_2_ at 37°C for 24h. After removing the medium, cells were incubated with 100 μL of fresh DMEM containing a suspension of promastigotes in the growth plateau phase at a rate of 1 cell per 10 parasites. After incubation under a 5 % CO_2_ atmosphere at 37°C for 24h (the time needed by the parasite to infect the macrophage), the culture medium was replaced with 100 μL of fresh DMEM with different concentrations of compounds as previously for a new incubation period of 48h. Controls were parasites alone in DMEM medium, axenic amastigotes, macrophages alone, infected macrophages and infected macrophages with different concentrations of miltefosine (Balaraman et al., 2015; Vásquez-Ocmín et al., 2018, 2021a).

#### Determination of IC_50_, CC_50_ and Selectivity Index for Plasmodium, Leishmania, and Trypanosma

After incubation, the plates were subjected to 3 freeze/thaw cycles to achieve complete cell lysis. The cell lysis suspension was diluted 1:1 except for Plasmodium plates that have been diluted 1:10 in lysis buffer (10 mM NaCl, 1 mM Tris HCl pH 8, 2.5 mM EDTA pH 8, 0.05 % SDS, 0.01 mg/mL proteinase K and 1X SYBR Green). SYBR Green incorporation in cell DNA amplification was determined using the Master epRealplex cycler^®^ (Eppendorf, France) and the following program to increase SYBR Green incorporation: 90°C for 1 min, decrease in temperature from 90°C to 10°C for 5 min with fluorescence reading at 10°C for 1 min and a new reading at 10°C for 2 min. Molecules were tested in duplicate. Compounds with different SYBR Green fluorescence values from duplicates were retested. The cytotoxicity of the compounds was expressed as IC_50_ (concentration of drug inhibiting the parasite growth by 50 %, comparatively to the controls), CC_50_ (Cytotoxic Concentration 50 %: concentration inhibiting macrophages growth by 50 %). IC_50_ and CC_50_ were calculated by nonlinear regression using ICEstimator 1.2 version (http://www.antimalarial-icestimator.net/MethodIntro.htm). Selectivity index (SI) for antiplasmodial and anti-*Trypanosoma* activities were calculated as the HUVEC’s CC_50_ divided to IC_50_ of *Plasmodium* 3D7 and *Trypanosoma*, and for *Leishmania* assays as the ratio of RAW 364.7 CC_50_ to intramacrophage amastigote IC_50_ value (Vásquez-Ocmín et al., 2018, 2021a).

#### In vitro evaluation on bloodstream form of Trypanosoma brucei gambiense

For each extract, twelve two-fold serial dilutions from 100 μg/mL to 0.049 μg/mL were performed in 96-well microplates in 100 μL HMI9 medium. Parasites were then added to each well to reach a final density of 4.104 cells/mL in 200 μL. Following 3 days of incubation at 37°C with 5 % CO_2_ in a water-saturated atmosphere, 20 μL of resazurin at 450 μM in water was added to each well for a final concentration of 40.9 nM to evaluate cell viability. Plates were then incubated for 6h at 37°C with 5 % CO_2_ in water-saturated atmosphere and the conversion of resazurin to resorufin was quantified by measuring the absorbance at 570 nm (resofurin) and 600 nm (resazurin) using the microplate reader Spark^®^ (Lyon, France) Pentamidine di-isethionate was used as the reference compound (Pomel et al., 2015; Vásquez-Ocmín et al., 2018, 2021a).

### Antimicrobial Assay

Minimal Inhibitory Concentration (MIC) determinations of crude extracts were carried out using the agar dilution method stipulated by the Clinical and Laboratory Standards Institute (CLSI, 2006). Antimicrobial activity was evaluated for the first time against a panel of 36 pathogenic and multi-drugresistant bacteria, which, in most cases, have been recently isolated from human infections. For comparison, reference strains from the American Type Culture Collection (ATCC) were included.

The inhibitory concentrations ranged between 0.075 and 1.2 mg/mL in five dilutions (1.2, 0.6, 0.3, 0.15 and 0.075 mg/mL); 0.075 mg/mL was considered a low enough concentration for a preliminary screening. Petri dishes were inoculated with strains (104 CFU, obtained by dilution in brain heart medium, BH) using a Steer’s replicator and were incubated at 37°C for 24 h. MIC was defined as the lowest concentration of extract without bacterial growth after incubation. The extracts with MIC ≤ 1.2 mg/mL were tested in triplicate at lower concentrations (mean absolute deviation is done for values: 1.2 ± 0.4; 0.6 ± 0.2; 0.3 ± 0.1; 0.15 ± 0.05; 0.075 ± 0.03). The standards (gentamicin, vancomycin, amoxicillin, amphotericin B, fluconazole, and sertaconazole) were tested in triplicate in 12 concentrations ranging from 0.03 to 64 mg/mL.

### Cytotoxic assay

Cytotoxicity was evaluated on RAW 264.7 macrophages for *Leishmania* assays and HUVECs for *Plasmodium* and *Trypanosoma*. The cells were seeded at a density of 50,000 cells per well in 100 μl of DMEM in a 96-well microtiter plate. After incubation in a 5 % CO_2_ at 37°C for 24h, the culture medium was replaced with 100 μl of fresh DMEM containing serial dilutions of the compounds tested. The concentrations of the compounds are the same as for intramacrophage *Leishmania* or *Plasmodium* assays. The plates were incubated for 48h at 37°C with 5 % CO_2_. Antiprotozoal assays have been performed only with compounds not demonstrating cytotoxicity (Vásquez-Ocmín et al., 2021a).

### Liquid chromatography and mass spectrometry data mining

#### HPLC-MS analyses

Extracts were analyzed by liquid chromatography performed on an Agilent 1260 series HPLC coupled to a 6530 QToF (Agilent Technologies). Chromatography separations were performed on a XSelect column C18, 2.1 x 75 mm – 2.5 μm (Waters). The mobile phase comprised water (0.1 % formic acid) (A) and acetonitrile (ACN) (B). A stepwise gradient method at a constant flow rate of 0.35 mL/min was applied as follows: 5–100% B (0–9.5 min), 100% B (4.5 min) and 4 min equilibration at 5% B. The mass spectrometer settings were: positive ESI mode, 50-3200 mass range calibration, and 2 GHz acquisition rate. Ionization source conditions were drying gas temperature 325 °C, drying gas flow rate 10 L/min, nebulizer 35 psig, fragmentor 150 V, and skimmer 65 V. Range of *m/z* was 200-1700. Purine C5H4N4 [M+H]^+^ ion (*m/z* 121.050873) and the hexakis-(1H,1H,3H-tetrafluoropropoxy)-phosphazene C18H18F24N3O6P3 [M+H]^+^ ion (*m/z* 922.009798) were used as internal lock masses. Full scans were acquired at a resolution of 11 000 (at m/z 922). MS-MS acquisitions were performed using three collision energies: 10, 20, and 40 eV. Three of the most intense ions (top 3) per cycle were selected. MS-MS acquisition parameters were defined as follows: *m/z* range 100-1200, default charge of 1, minimum intensity of 5000 counts, rate/time = 3 spectra/s, isolation width: Narrow (1.3 u).

#### Data processing

For untargeted metabolomics, the LC-MS data were processed according to the MSCleanR workflow (Fraisier-Vannier et al., 2020; Vásquez-Ocmín et al., 2021b). Briefly, a batch in positive ionization (PI) was processed with MS-DIAL version 4.90 (Tsugawa et al., 2015). MS1 and MS2 tolerances were set to 0.01 and 0.05 Da, respectively, in centroid mode for each dataset. Peaks were aligned on a QC reference with an RT tolerance of 0.2 min, a mass tolerance of 0.015 Da, and a minimum peak height detection at 1 × 10^5^. MS-DIAL data was deconvoluted together with MS-CleanR by selecting all filters with a minimum blank ratio set to 0.8 and a maximum relative standard deviation (RSD) set to 40. The maximum mass difference for feature relationship detection was set to 0.005 Da, and the maximum RT difference was set to 0.025 min. Pearson correlation links were used with a correlation ≥ 0.8 and a p-value significance threshold of 0.05. Two peaks were kept in each cluster for further database requests and the kept features were annotated with MS-FINDER version 3.52 (Tsugawa et al., 2016). The MS1 and MS2 tolerances were set to 5 and 15 ppm, respectively. The formula finder was exclusively based on C, H, O, and N atoms. Three levels of compounds annotation from several sets of data were carried out. Metabolite annotation at level 1 using MS-DIAL: i) the experimental LC-MS/MS data of 500 compounds (retention time, exact mass and fragmentation) were used as references; ii) the Mass spectral records from MS-DIAL, MONA (MassBank of North America), and GNPS (Global Natural Product Social Molecular Networking) databases were used for spectral match applying a dot product score cut-off of 800. Metabolite annotation at level 2 was prioritized according to: i) a search would be made using MSFinder for a match with compounds identified in the literature for the *Piper* genus (genus level) and Piperaceae family (family level) (Dictionary of Natural Products version 28.2, CRC Press) based on exact mass and *in silico* fragmentation. For metabolite annotation level 3, a search was made using MS-FINDER for a match with natural compounds included in their databases embedded (PlantCyc, ChEBI, NANPDB, COCONUT, and KNApSAcK) (generic level) (Vásquez-Ocmín et al., 2022b).

Mass spectra similarity networking was carried out from PI mode using MetGem (Olivon et al., 2018) on the final annotated .msp file and metadata file for PI obtained with MSCleanR. Values used were MS2 m/z tolerance = 0.02 Da, minimum matched peaks = 4 and minimal cosine score value = 0.7. The visualization of the molecular network (MN) was performed on Cytoscape version 3.9.1 (Shannon et al., 2003). The list of compound mass, retention time, row ID and peak height was exported in CSV format. Then, this format was used for the statistical data analysis.

#### Statistical analyses

The final annotated untargeted metabolome feature matrix and biological assay results datasets were analyzed using the R package MixOmics (http://mixomics.org/), which is dedicated to the integrative analysis of ‘omics’ data (Le Cao et al., 2016). Biological assays results expressed in IC_50_ were transformed in pIC_50_ (-log10 [IC_50_]) for further statistical correlation analysis. MIC values were log transformed. The LC-MS dataset (*m/z* × RT × peak area) was normalized to total ion chromatogram and scaled to unit variance. A vertical integration approach has been afforded to leverage on multiplexed measurements for the same extract (González et al., 2008). To highlight the overall correlation between chemical fingerprints and biological assays results, a regularized Canonical Correlation Analysis (rCCA) was done using a cross validation approach for tuning regularization parameters. The correlation structure of the two-block data matrices was displayed on clustered image map (CIM) and relevance network using 0.3 as cutoff correlation value.

## RESULTS

### Ethnopharmacological survey

Our set of 19 samples was collected in three Amazonian regions of Peru, where the use of these species of “*Cordoncillos*” in traditional medicine is widespread. All information about the collected samples are presented in **supp info 1**. Among the plants studied in this work, 10 (52.6 %) had not been subject to chemical analysis so far. In our previous work (Vásquez-Ocmín et al., 2021a), *Piper casapiense, P. strigosum* and *P. pseudoarboreum* showed promising antiprotozoal activity. Thus, these plants were re-collected in other areas of the Loreto region.

### Anti-infectious activity

#### Antiprotozoal activity

The IC_50_ values for antiplasmodial, antileishmanial and antitrypanosomal activities of each extract are shown in **Table 1**. We considered an extract active and worthy to be further studied when its IC_50_ value was below 10 μg/mL. Four extracts were active in this case regarding the 3D7 *P. falciparum* chloroquine-sensitive strain assay: the hexane and methylene chloride extracts of *P. crassinervium*, the methanol extract of *P. stellipilum*, and the methylene chloride extract of *P. xanthostachyum*. Only the methylene chloride extracts of *P. crassinervium*, and *P. oblongum* were active on three strains of *Leishmania* (both axenic and intra-macrophagic amastigotes). For the other active extracts, six were not active on intra-macrophagic forms: hexane and methylene chloride of *P. heterophyllum*, hexane, methylene chloride and water extracts of *P. sancti-felicis* (*L. amazonensis*) and a hexane extract of *P. crassinervium (L. donovani*). Hexane extract of *P*. “*cordatomentosa*” was active on all the strains, both axenic and intramacrophagic amastigotes, except on axenic amastigote forms of *L. donovani*. Five extracts were active only on *L. amazonenesis*, and *L. braziliensis* (both axenic and intramacrophagic amastigotes): methylene chloride extracts of *P. calvescentinerve*, *Piper crassinervium*, and *P. oblongum*, and the methylene chloride and hexane extracts of *Piper “cordatomentosa”, and Piper crassinervium*. Five extracts were active only on *L. donovani*, and *L. braziliensis* (both axenic and intramacrophagic amastigotes), the methylene chloride extract of *P. crassinervium*, and the hexane and methylene chloride extracts of *P. heterophyllum*, and *P. oblongum*. Hexane extract of *P. glabribaccum* was not active on axenic amastigote forms of *L. donovani*. Two extracts were active on *L. donovani*, and *L. amazonenesis*, both forms, methylene chloride extracts of *Piper crassinervium*, and *P. oblongum*. Selective activity on *L. braziliensis* were observed for twelve extracts, the hexane and methylene chloride extracts *Piper sancti-felicis*, *Piper stellipilum*, and *P. xanthostachyum*; hexane and methylene chloride extracts of *P. stellipilum*, and hexane and methylene chloride extracts of *P. calvescentinerve* and of *P. divaricatum*. None of the extracts showed selective activity on *L. donovani* or *L. amazonensis*.

**Table 1.**
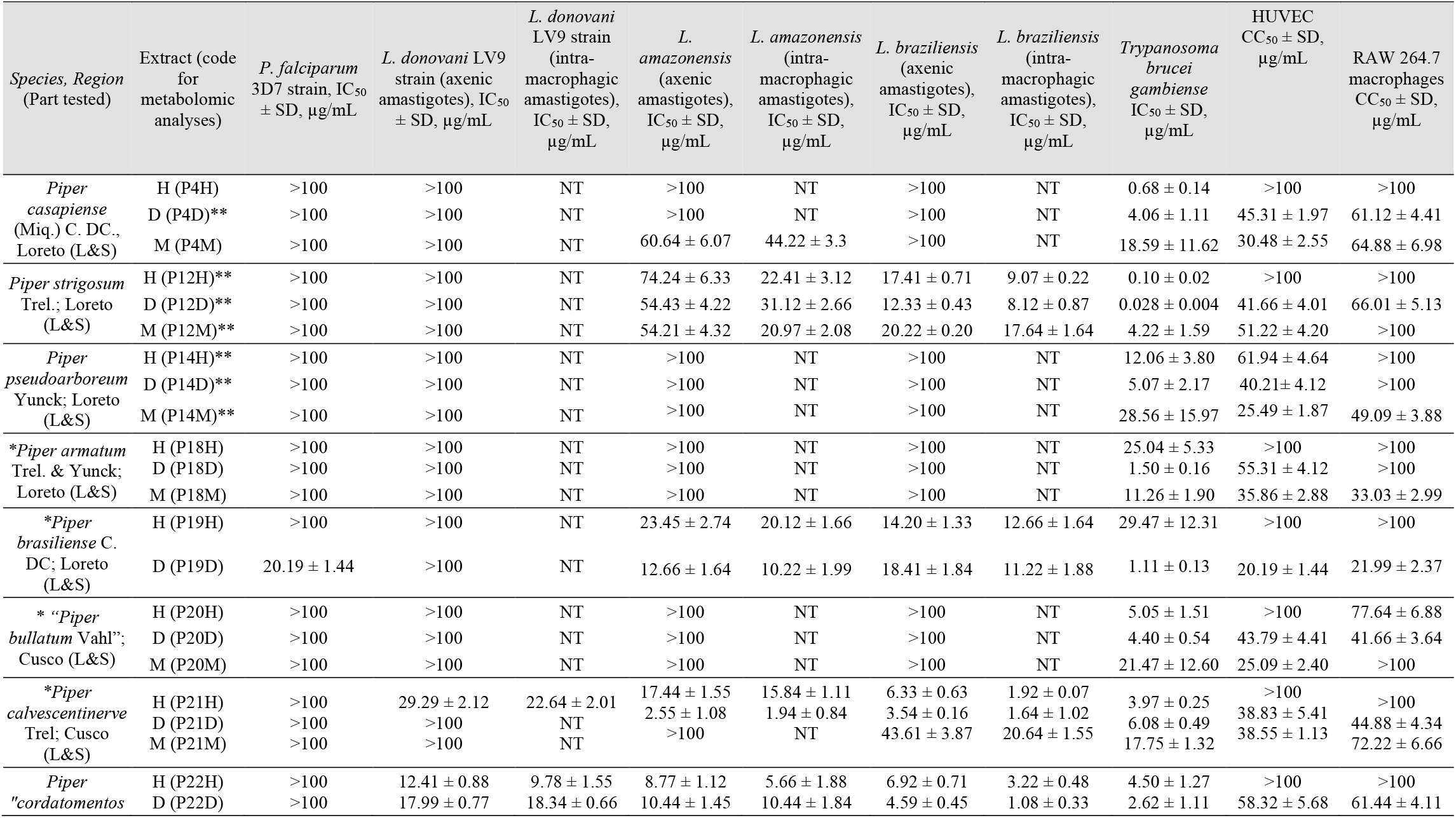
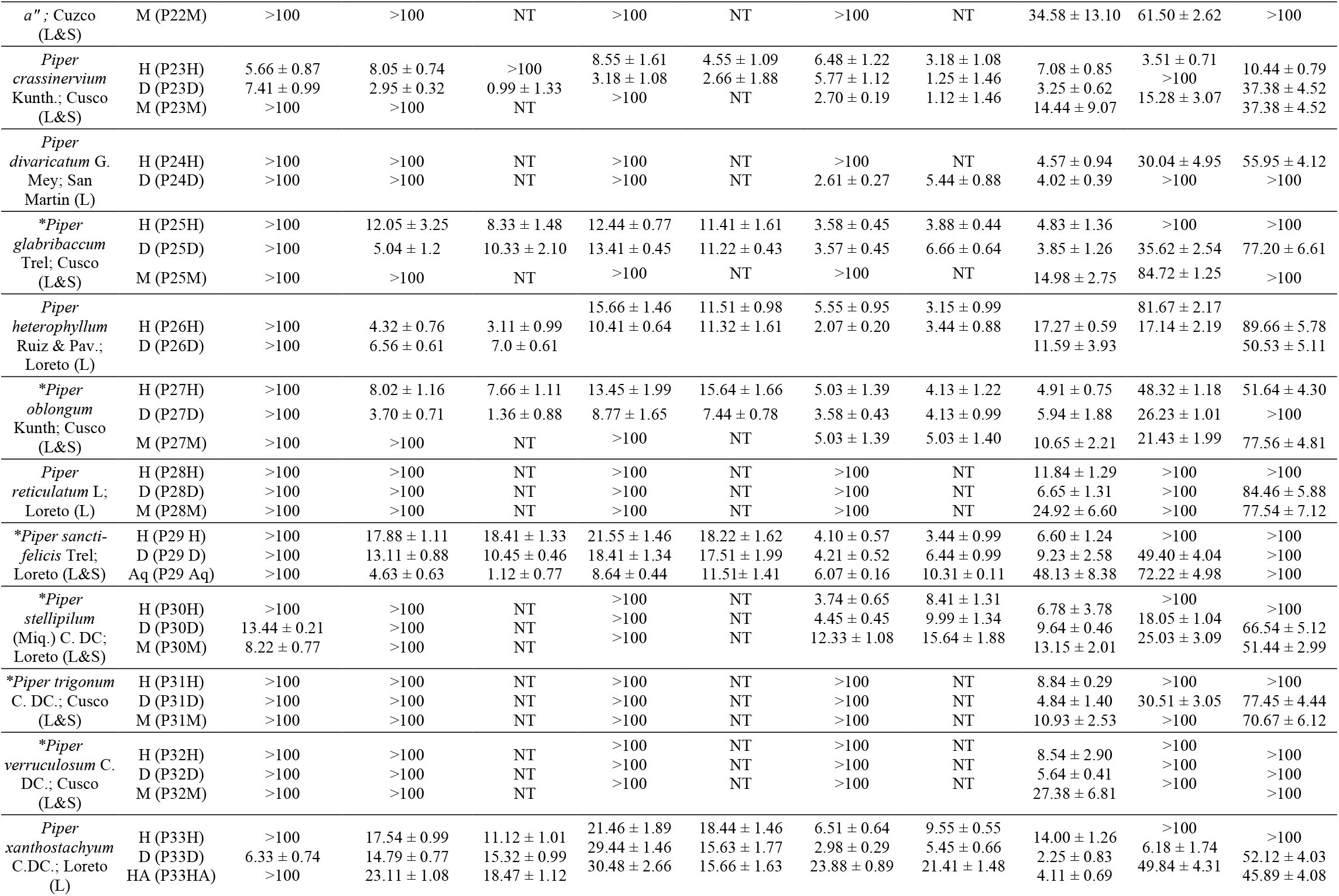
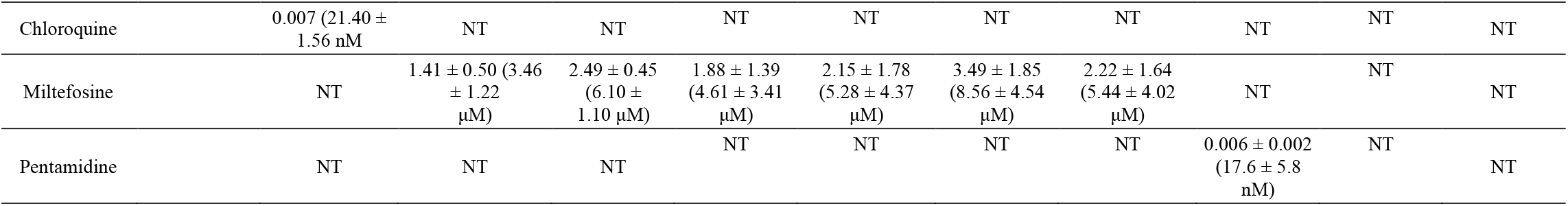
*In vitro* antiprotozoal and cytotoxicity activities of *Piper* extracts. M = methanol; H = hexane; D = methylene chloride (D stands for dichloromethane); HA = hydroalcoholic; Aq = aqueous; NT = not tested. *Species showing no reference in phytochemical literature on Web of Science and PubMed. **Extracts previously tested (Vásquez-Ocmín et al., 2021a) (the plants in this work come from different collection site). L = leaves; L&S = Leaves and stems; AP= aerial parts; “” = species with temporary botanical name (these species need more taxonomical studies).

Selectivity index (SI) values for antiprotozoal activities (**Table 2**) were ranked as low >10<50, high >51<99, and very high >100. For antimalarial activity, the only active extract displaying a low SI (13.50) was the methylene chloride extract of *P. crassinervium*. For the antileishmanial activity, methylene chloride extract of *P. crassinervium* and hexane extracts of *Piper glabribaccum* and *P. heterophyllum* showed a low SI on *L. donovani* with 37.76, 12.0, and 28.82 respectively; methylene chloride and aqueous extracts of *P. oblongum* and *P. sancti-felicis* showed high SI with 73.53 and 89.29 respectively. For *L. amazonenesis*, three extracts presented a low SI, methylene chloride extracts of*P. “cordatomentosa”*, *P. crassinervium*, and *P. oblongum* with 13.39, 14.05, and 13.44 respectively. For *L. braziliensis*, 17 extracts displayed low SI, the hexane extracts of *P. strigosum*, *P. “cordatomentosa”*, *P. glabribaccum*, *P. heterophyllum*, *P. oblongum*, *P. sancti-felicis*, *P. stellipilum*, and *P. xanthostachyum* (11.02, 31.06, 25.77, 28.46, 12.50, 29.07, 11.89, 10.47), methylene chloride extracts of *Piper calvescentinerve*, *P. crassinervium*, *P. divaricatum*, *P. glabribaccum*, *P. heterophyllum, P. oblongum*, and *P. sancti-felicis* (27.37, 29.90, 18.38, 11.59, 14.69, 24.21, 15.53, respectively); methanol extracts of *Piper crassinervium*, and *P. oblongum* (33.36 and 15.42). Only the hexane and methylene chloride extracts of *P. calvescentinerve* and *P. “cordatomentosa”* presented a high SI with 52.08 and 56.89, respectively.

**Table 2.** Selectivity index of *Piper* extracts calculated with CC_50_ on HUVEC cells for *P. falciparum* and *T. b. gambiense*, and CC_50_ on RAW 264.7 cells for *Leishmania* strains. M = methanol; H = hexane; D = methylene chloride (D stands for dichloromethane); Aq = aqueous; ND = not determined. *Species with no phytochemical reference in Web of Science and PubMed. **Extracts previously tested (Vásquez-Ocmín et al., 2021a) (the plants come from different collection site). L = leaves; L&S = Leaves and stems. ““ = species with a temporary botanical name (these species need further taxonomic study).

| Species | Extract (code for metabolomic analyses) | SI on *P. falciparum* 3D7 strain, $CC_{50}/IC_{50}$ | SI on *L. donovani* (intra-macrophagic amastigotes), $CC_{50}/IC_{50}$ | SI on *L. amazonensis* (intra-macrophagic amastigotes), $CC_{50}/IC_{50}$ | SI on *L. braziliensis* (intra-macrophagic amastigotes), $CC_{50}/IC_{50}$ | SI on *T. b. gambiense*, $CC_{50}/IC_{50}$ |
| --- | --- | --- | --- | --- | --- | --- |
| *Piper casapiense* | H (P4H) | $\leq 1$ | ND | ND | ND | 147.06 |
|  | D (P4D)** | 0.45 | ND | ND | ND | 11.16 |
|  | M (P4M) | 0.30 | ND | 1.46 | ND | 1.64 |
| *Piper strigosum* | H (P12H)** | $\leq 1$ | ND | 4.46 | 11.02 | 1000 |
|  | D (P12D)** | 0.42 | ND | 2.12 | 8.13 | 1487.86 |
|  | M (P12M)** | 0.51 | ND | 4.77 | 5.67 | 12.14 |
| *Piper pseudoarboreum* | H (P14H)** | 0.62 | ND | ND | ND | 5.14 |
|  | D (P14D)** | 0.40 | ND | ND | ND | 7.93 |
|  | M (P14M)** | 0.25 | ND | ND | ND | 0.89 |
| *\*Piper armatum* | H (P18H) | ≤ 1 | ND | ND | ND | 3.99 |
|  | D (P18D) | 0.55 | ND | ND | ND | 36.87 |
|  | M (P18M) | 0.36 | ND | ND | ND | 11.26 |
| *\*Piper brasiliense* | H (P19H) | ≤ 1 | ND | 4.97 | 7.90 | 3.39 |
|  | D (P19D) | 1.00 | ND | 2.15 | 1.96 | 18.19 |
| *\*“Piper bullatum”* | H (P20H) | ≤ 1 | ND | ND | ND | 19.80 |
|  | D (P20D) | 0.44 | ND | ND | ND | 9.95 |
|  | M (P20M) | 0.25 | ND | ND | ND | 1.17 |
| *Piper calvescentinerve* | H (P21H) | ≤ 1 | 4.42 | 6.31 | 52.08 | 25.19 |
|  | D (P21D) | 0.39 | ND | 2.31 | 27.37 | 6.39 |
|  | M (P21M) | 0.39 | ND | ND | ND | 2.17 |
| *“Piper cordatomentosa”* | H (P22H) | ≤ 1 | 10.22 | 1.77 | 31.06 | 22.22 |
|  | D (P22D) | 0.58 | 3.35 | 13.39 | 56.89 | 22.26 |
|  | M (P22M) | 0.62 | ND | ND | ND | 1.79 |
| *Piper crassinervium Kunth* | H (P23H) | 0.62 | 0.10 | 2.99 | 3.28 | 0.50 |
|  | D (P23D) | 13.50 | 37.76 | 14.05 | 29.90 | 30.77 |
|  | M (P23M) | 0.15 | ND | ND | 33.36 | 10.58 |
| *Piper divaricatum* | H (P24H) | 0.30 | ND | ND | ND | 6.57 |
|  | D (P24D) | ≤ 1 | ND | ND | 18.38 | 24.88 |
| *\*Piper glabribaccum* | H (P25H) | ≤ 1 | 12.00 | 8.76 | 25.77 | 20.70 |
|  | D (P25D) | 0.36 | 7.47 | 0.15 | 11.59 | 9.25 |
|  | M (P25M) | 0.85 | ND | ND | ND | 5.66 |
| *Piper heterophyllum* | H (P26H) | 0.82 | 28.82 | 7.79 | 28.46 | 4.73 |
|  | D (P26D) | 0.17 | 7.22 | 4.46 | 14.69 | 1.48 |
| *\*Piper oblongum* | H (P27H) | 0.48 | 6.74 | 3.30 | 12.50 | 9.84 |
|  | D (P27D) | 0.26 | 73.53 | 13.44 | 24.21 | 4.42 |
|  | M (P27M) | 0.21 | ND | ND | 15.42 | 2.01 |
| *Piper reticulatum* | H (P28H) | ≤ 1 | ND | ND | ND | 8.45 |
|  | D (P28D) | ≤ 1 | ND | ND | ND | 15.04 |
|  | M (P28M) | ≤ 1 | ND | ND | ND | 4.01 |
| *\*Piper sancti-felicis* | H (P29 H) | ≤ 1 | 5.13 | 5.49 | 29.07 | 15.15 |
|  | D (P29 D) | 0.49 | 9.57 | 5.71 | 15.53 | 5.35 |
|  | Aq (P29 Aq) | 0.72 | 89.29 | 8.69 | 9.70 | 1.50 |
| *\*Piper stellipilum* | H (P30H) | ≤ 1 | ND | ND | 11.89 | 14.75 |
|  | D (P30D) | 1.34 | ND | ND | 6.66 | 1.87 |
|  | M (P30M) | 3.04 | ND | ND | 3.29 | 1.90 |
| *\*Piper trigonum* | H (P31H) | ≤ 1 | ND | ND | ND | 11.31 |
|  | D (P31D) | 0.31 | ND | ND | ND | 6.30 |
|  | M (P31M) | ≤ 1 | ND | ND | ND | 9.15 |
| *Piper verruculosum* | H (P32H) | ≤ 1 | ND | ND | ND | 11.70 |
|  | D (P32D) | ≤ 1 | ND | ND | ND | 17.73 |
|  | M (P32M) | ≤ 1 | ND | ND | ND | 3.65 |
| *Piper xanthostachyum* | H (P33H) | ≤ 1 | 8.99 | 5.42 | 10.47 | 7.14 |
|  | D (P33D) | 0.98 | 3.40 | 3.33 | 9.56 | 2.75 |
|  | HA (P33HA) | 0.50 | 2.48 | 2.93 | 2.14 | 12.13 |

At least one extract of all nineteen plants, except *P. heterophyllum*, was active on *T. b. gambiense* at IC_50_ ≤ 10 μg/mL (**Table 1**). The only two plants that displayed activity in all extracts were *P. strigosum* and *P. divaricatum*. Among all active extracts, five had very good activity with a IC_50_ ≤ 1 μg/mL, hexane and methylene chloride extracts of *P. strigosum*, hexane extract of *P. casapiense*, and methylene chloride extracts of *P. armatum*, and *P. brasiliense*. Especially, the methylene chloride extract from *Piper strigosum* was very active with an IC_50_ at 0.028 μg/mL. Three extracts presented a high SI, hexane extract of *P. casapiense* with 147.06 and the hexane and methylene chloride extracts of *P. strigosum* with 1000 and 1487.86, respectively. Twenty extracts displayed a lower yet interesting SI between 10.58 and 36.87.

#### Antimicrobial activity

Antimicrobial assay results are presented in Table 3 (17 Gram-positive bacteria, 13 Gram-negative bacteria, 2 yeasts). Extracts were more active on Gram-positive bacteria and yeast. The IC_50_ cut-off values were set at ≤ 0.5 mg/mL for good activity and ≤ 0.09 mg/mL for very good activity. Among the very active extracts, methylene chloride extract of *P. strigosum* is the most representative, being very active on all Gram-positive strains [(excepted: *Staphylococcus warneri* (T26A1)], only on two Gramnegative strains *[(Stenotrophomonas maltophilia* (21170) and *Burkholderia cepacia* (13003)], and on the two *Candida albicans* strains. This extract was also quite active on *Streptococcus pyogenes* (16135). Other extracts with good activity on Gram-positive bacteria can be pointed out: methylene chloride extracts of *P. xantochyma* and *P. brasiliense*, and hydroalcoholic extract of *P. xantochyma*. Extracts with very good activity on Gram-negative bacteria are: methylene chloride extracts of *P. xantochyma* and *P. divaricatum* on *Burkholderia cepacia* (13003) and *Pseudomonas aeruginosa* (ATCC 27583), respectively. Extracts with very good activity on *Candida* strains were methylene chloride extracts of *P. pseudoarboreum*, *P. divaricatum*, and *P. heterophyllum* and hexane extract of *P. sancti-felicis*. Extracts with good activity on Gram-positive bacteria and *Candida* were principally methylene chloride of *P. brasiliense* and a hydroalcoholic extract of *P. xantochyma*.

**Table 3.**
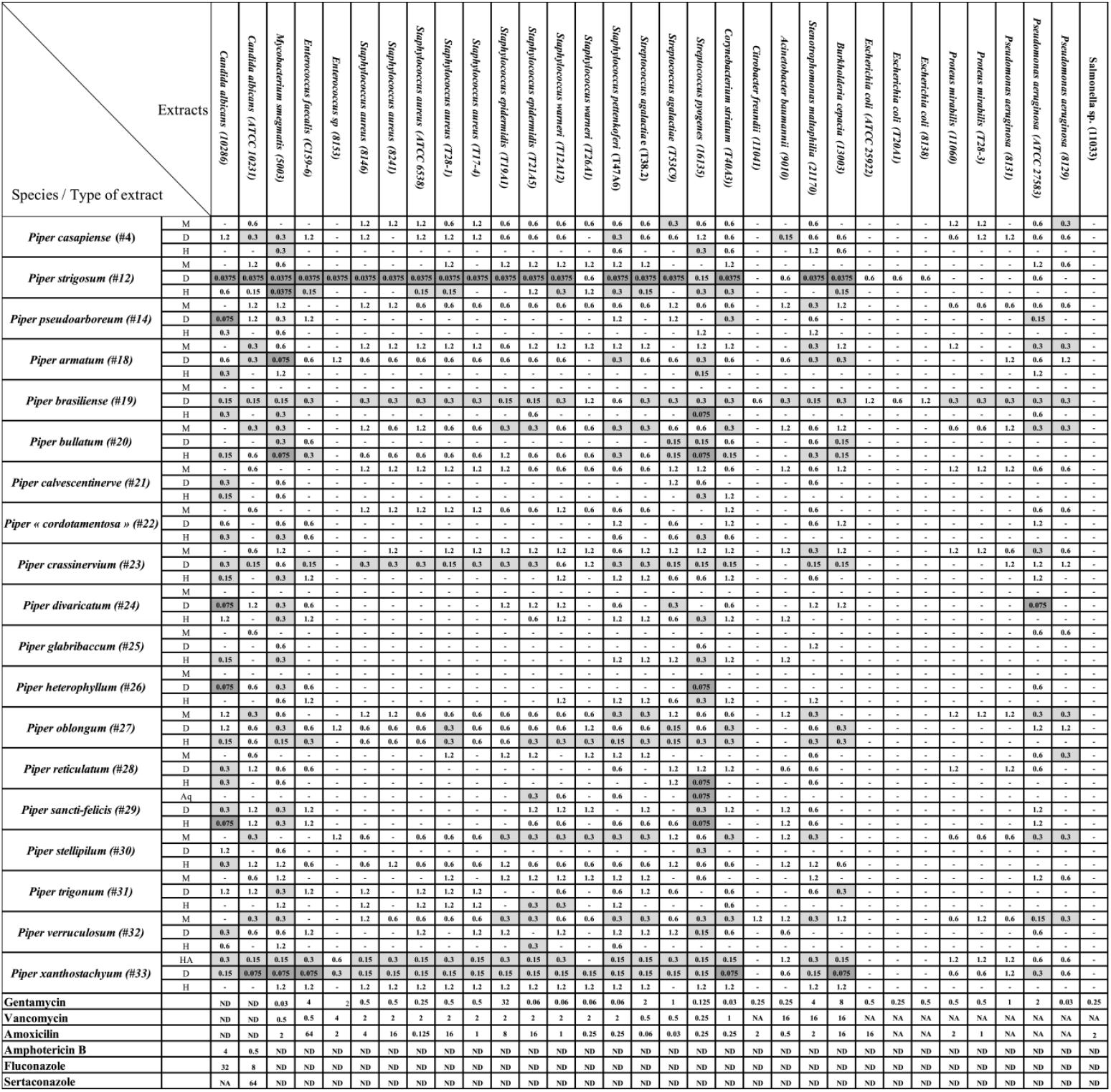
*In vitro* antimicrobial activity of *Piper* extracts (minimal inhibitory concentration MIC), in mg/mL). ND = not determinate, NA = not active.

#### LCMS data mining and statistical correlation analysis

After the application MS-CleanR workflow to the LCMS data, we obtained 123 unique metabolite features (*m/z* × RT × Peak area). Among these, 56 compounds were annotated at the genus level (45.53 %), 6 compounds at the family level (4.88 %), and 55 compounds at the generic level (44.72 %), leaving 6 compounds unannotated (4.88 %). The MSMS fragmentation pathway of these compounds was used to establish a molecular network (**Figure 1**). Using NPClassifire and ClassyFire (Djoumbou Feunang et al., 2016; Kim et al., 2021), major phytochemical classes and subclasses of compounds were identified. The main classes of compounds identified were alkaloids, amino acids, fatty acids, polyketides, shikimates, phenylpropanoids, and terpenoids (metadata in **supp info 2**).

**Figure 1.**
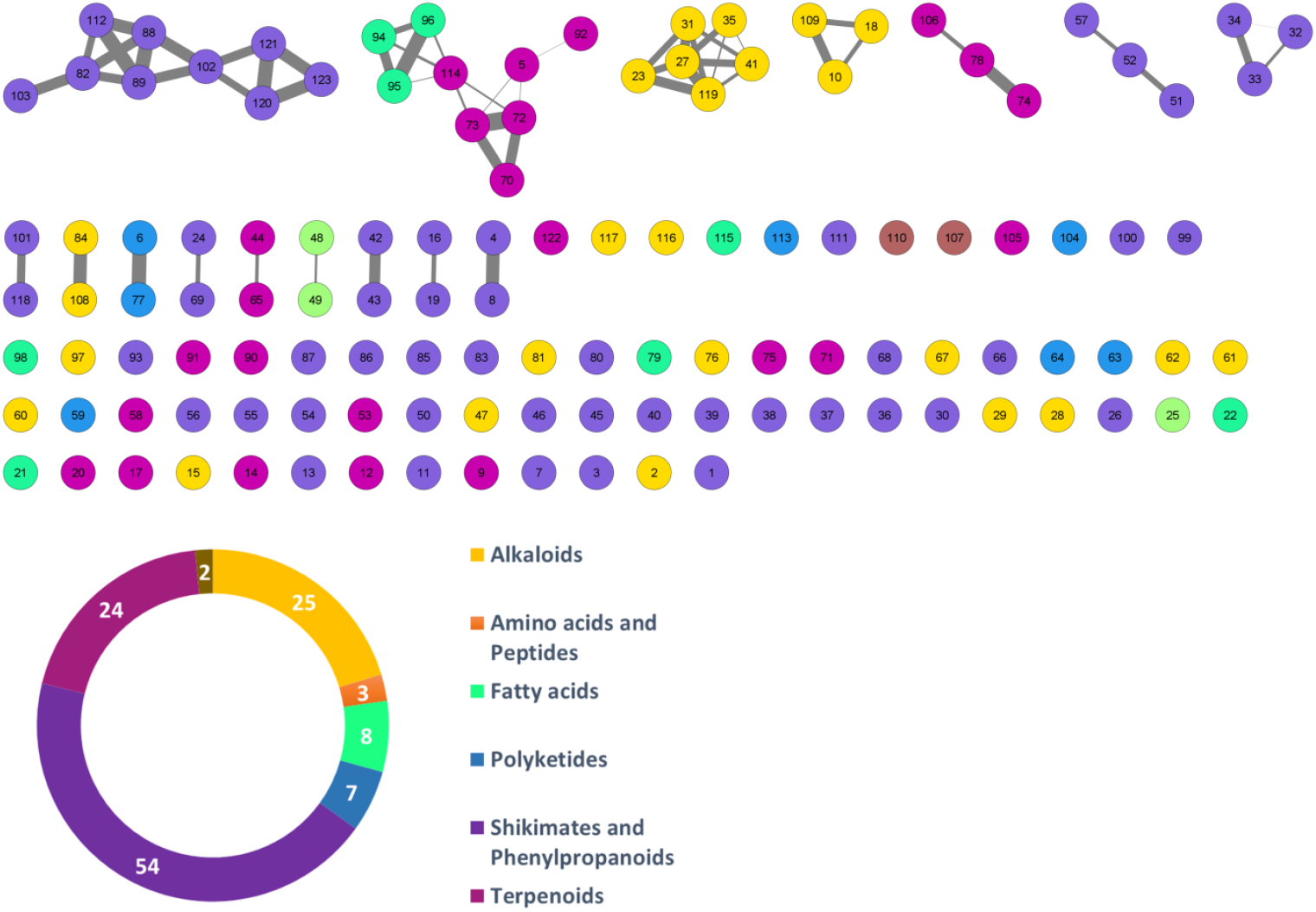
Molecular networking of putative compounds. A feature ID is allocated in each node. The colors in the network and pie chart relate to the phytochemical classes. The number of features for each class is also shown in the pie chart.

A heatmap of correlation analysis (Clustered Image Map, CIM) is shown in **Figure 2-A**. For this analysis, we compiled all *in vitro* assay results (30 bacterial strains, 2 fungal strains, 4 parasitic strains and 2 cell lines, **Tables 1** and **3**) *versus* the LCMS features. This figure results from a setting of the threshold of correlation at 0.3 (the heatmap is shown *in extenso* in **supp info 3**). Two groups of features responsible for activities can be distinguished. **Group 1** is composed of 11 features (12, 21, 23, 27, 31, 35, 41, 45, 87, 95, and 119) having activity on 21 bacteria, one fungus (*C. albicans* ATCC 10231), and one parasite (*T. b. gambiense*). Activity on bacteria was mostly concentrated on Gram-positive strains [(except for *Burkholderia cepacia* (13003), *Stenotrophomonas maltophilia* (21170), and the three strains of *Escherichia coli)]*. **Group 2** is composed of 9 features (4, 6, 7, 16, 53, 63, 65, 77, and 117). This group was selective on *Leishmania* (all strains, both axenic and intramacrophagic). We also performed a relevance network (RN) analysis on the same datasets to verify and complete the CIM results (**Figure 2-B**). RN showed two main clusters of features linked to the activity, the same as in the CIM. The LC-MSMS molecular network (**Figure 1**) allowed to determine that six features identified in the **Group 1** belong to the alkaloids class, while others are mostly self-loops (*i. e*. compounds not having structurally-related analogs in the extract) belonging to varied phytochemical classes. In the case of *Leishmania*-specific compounds (**Group 2**), they mostly belong to the polyketides or shikimates and phenylpropanoids classes, and are found only as self-loops or only related to one compound and not correlated to any activity (*i.e*. features 16 and 4).

**Figure 2.**
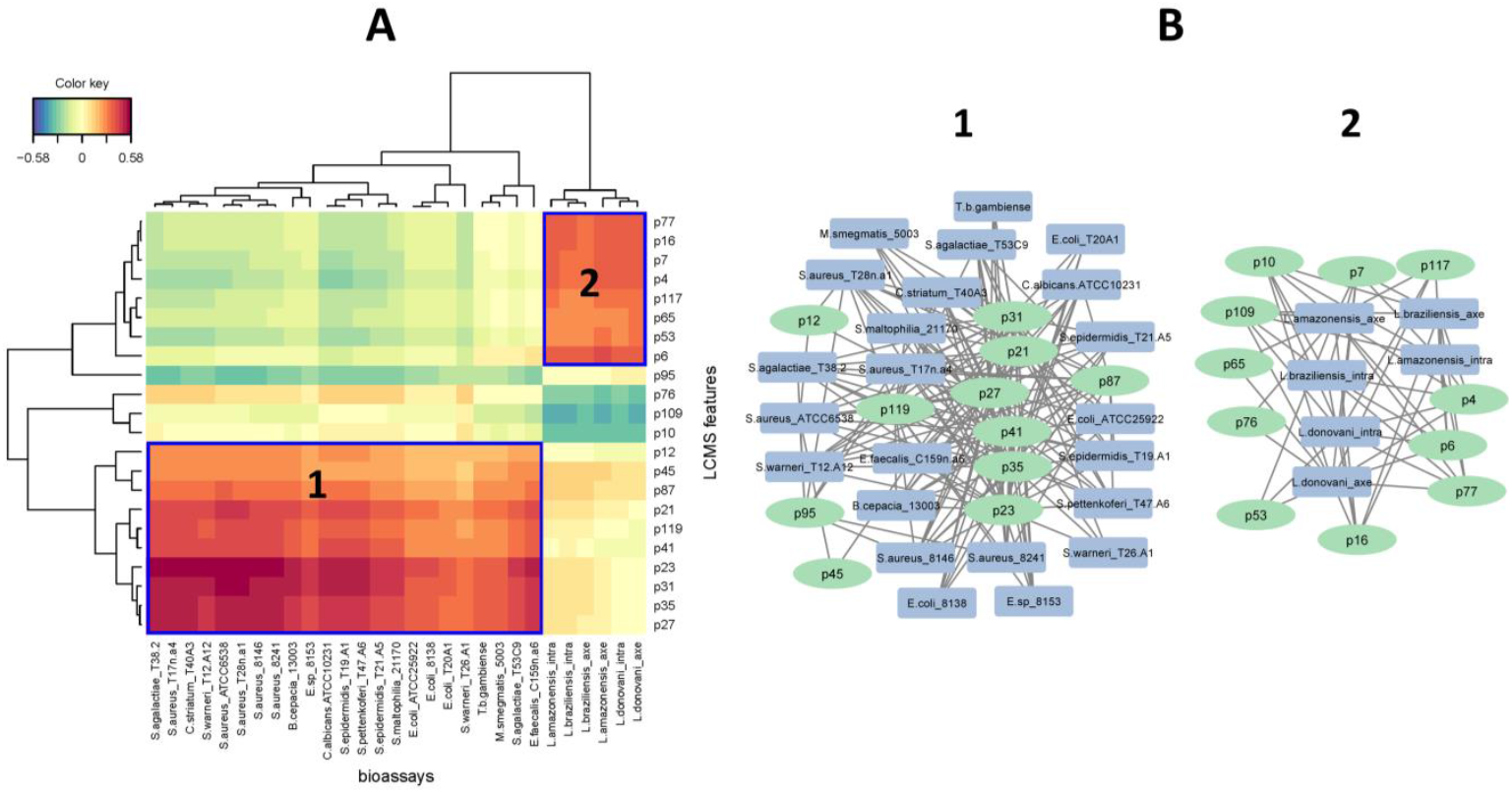
A) Clustered Image Map (CIM) performed on the two-blocks of data sets. The plot depicts the correlation between X = LCMS features (p = peak) data and Y = bioassays. Variables were selected across two dimensions and clustered with a complete Euclidean distance method. Blue squares represent the two main groups of features identified by activities B) The two main clusters of interaction represented by the relevance network (RN). The two analyses were tuned to a threshold of interaction of 0.3.

**Table 4.**
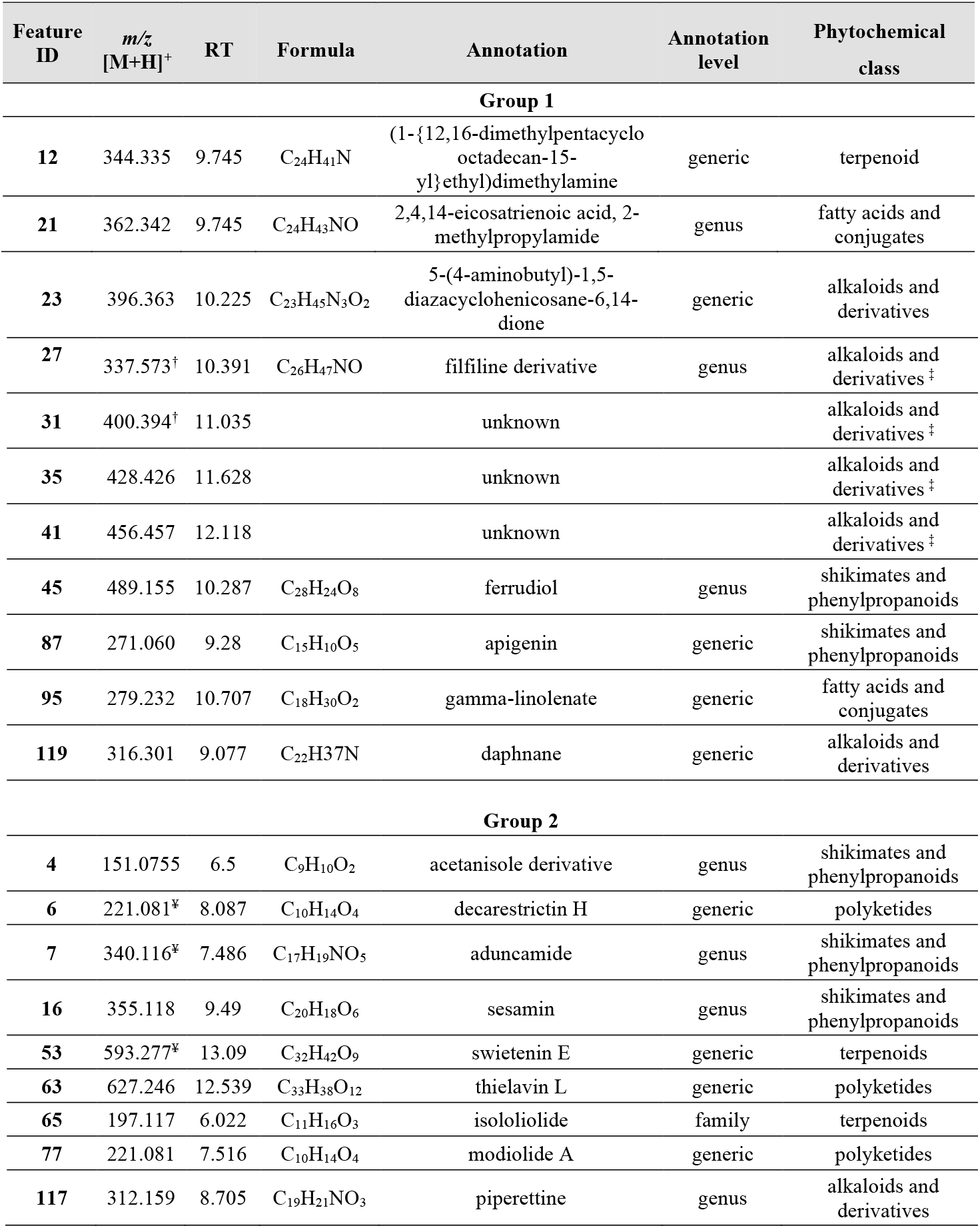
Features identified as responsible for bioactivity. ^†^ = [M+H-H_2_O]^+^; ^¥^= [M+Na]^+^; ^‡^= based on molecular networking.

**Figure 3** shows a heatmap depicting how the features are distributed among the extracts. Features of **Group 1** are mainly distributed in all extracts of *P. strigosum* (features 12, 21, 23, 31, 41, 27 and 35) and *P. xanthostachyum* (features 45 and 87). Features of **Group 2** are found in the hexane and methylene chloride extracts of *P. sancti-felicis* and hexane extracts of *P. calvescentinerve*, *P. cordatomentosa* and *P. crassinervium* (features 6, 7, 16 and 77), methylene chloride extracts of *P. pseudoarboreum*, *P. calvescentinerve, P. divaricatum*, *P. glabribaccum*, *P. bullatum*, *P. heterophyllum*, *P. reticulatum*, *P. oblongum, P. trigonum*, *P. crassinervium* and *P. cordatomentosa*, and the aqueous extract of *P. sancti-felicis* (features 4, 53, 63, 65 and 117). Feature 95 is found in the methanol and hexane extracts of *P. reticulatum* and the hexane extracts of *P. pseudoarboreum*, *P armatum*, *P. glabribaccum*, *P. oblongum* and *P. verruculosum*.

**Figure 3.**
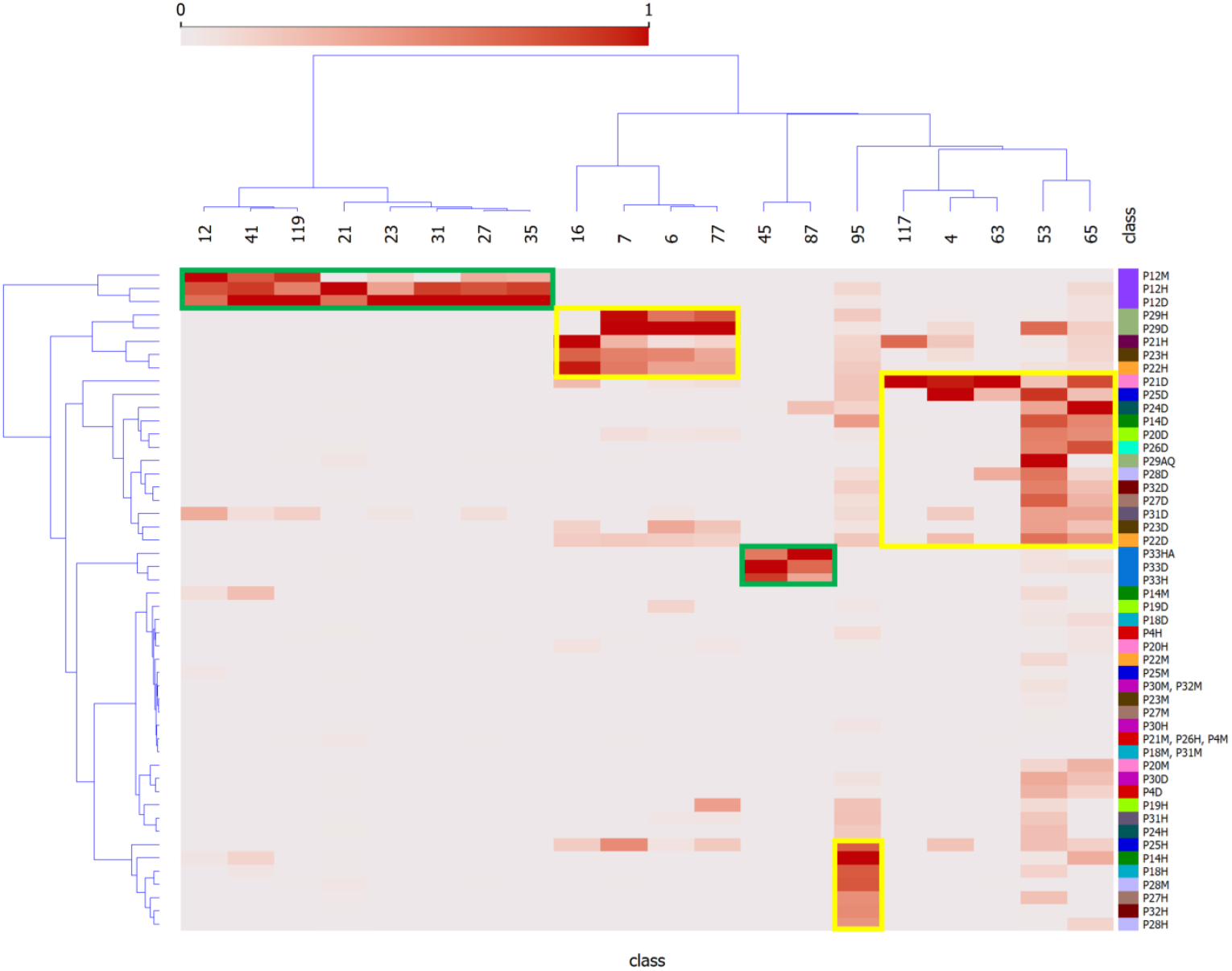
Heatmap of correlation between plant extracts and presumed bioactive peak features. Y = *Piper* extracts, and X = features of LC-MSMS analysis as shown in Figure 2A-B. Green and yellow squares represent the distribution of the bioactive features identified for the Groups 1 and 2 in the *Piper* extracts.

## DISCUSSION

Plants from the *Piper* genus are currently used by the Amerindian communities in Peru in different preparations (infusion of different parts, juice directly ingested, cataplasms made of crushed leaves or stems, etc.) to treat different diseases (Odonne et al., 2009, 2011; Vásquez-Ocmín et al., 2021a). For the present study, we conducted ethnopharmacological surveys in Loreto and Cusco, Peruvian Amazonian regions heavily impacted by protozoal diseases. Indeed, Loreto has always been the department with the greatest number of malaria cases (principally *P. vivax*), mostly affecting children from birth to age 11 (84.5 % in 2020) (Ministerio de Salud del Perú, 2021). Leishmaniasis occurrence in Peru is principally due to cutaneous (*L. amazonesis*, 17 892 cases over the last five years) and mucosal forms (*L. braziliensis*, *L. peruviana*, *L. lainsoni*, *L. guyanensis*, 1 642 cases in the last five years). To date, in 2022 70% of the cases of both forms were concentrated in 7 departments: Madre de Dios, Cusco (Echarate and Kosñipata districts), Piura, Junín, Loreto, Cajamarca and San Martín (Ministerio de Salud del Perú, 2022). In Peru, plant-based treatments are very common and even recommended by the health personnel. It is estimated that approximately 5000 species are used in the traditional medicine (Vásquez-Ocmín et al., 2018) among which the *Piper* species studied in this work represent 0.34 %. We targeted the ethnopharmacological survey on *“Cordoncillos”* (little cords), a word used to describe pepper vines in Amazonia because the presence of nodes on the stems reminds us of a *nudo de soga* (node of cord). While this terminology would initially direct the sample collection, we took great care of properly identifying the species upon collection. In two cases, we could not fully ascertain the species and further taxonomic examination is underway.

Chemical profiling of natural extracts can be readily obtained, but for metabolomic purposes, data need to be ‘preprocessed’ (peak picking, alignment, clustering, integration and normalization) to obtain a clean dataset able to provide meaningful information upon statistical analysis. The large abundance and complexity of variables are also challenging and require to apply an appropriate statistical method to answer a specific biological question (Hervé et al., 2018). Few metabolomic studies on *Piper* species using NMR (Yamaguchi et al., 2011; Uckele et al., 2021) or LCMS have been reported so far (Vásquez-Ocmín et al., 2021a). The *Piper* genus is a good model to apply our framework for two reasons: i) *Piper* species are particularly abundant in Peru (~260 according to www.tropicos.org), many of them used in traditional medicine; ii) many studies, including numerous phytochemical ones, were carried out on this genus (~9000 results found in www.webofscience.com for *“Piped”*, consulted on July 1^st^, 2022), allowing a good metabolome coverage.

We propose in this work a computational statistical analysis to integrate two heterodimensional datasets: the features (compounds) obtained by LC-MSMS analysis of *Piper* extracts (**supp info 2**), and the *in vitro* biological activity of these extracts on parasites, bacteria and mammalian cells (**Tables 1** and **3**), with the aim to outline most active compounds. To leverage on this multiplexed approach, we applied a *r*egularized *C*anonical *C*orrelation *A*nalysis (rCCA). This method aims to extract correlated information by maximizing their correlation via canonical variates between two datasets acquired on the same samples (vertical integration). The results of rCCA model were displayed using Clustered Image Maps (CIM) to highlight the correlation level of variables from all datasets, ordered through unsupervised clustering on both samples and compounds simultaneously (González et al., 2012). Another approach for displaying net-like correlation structures between two data sets is to use Relevance Networks (RN). This method generates a graph where nodes represent variables, and edges represent variable associations (Butte et al., 2000). Given an appropriate threshold, the RN highlights main correlation between both datasets (**Figure 2-B**). To our knowledge, the implementation of this type of metabolomics framework aimed at identifying bioactive compounds has not been used so far. However, a study has been recently published using rCCA to combine multiparametric analysis of adjuvanticity *in vivo* with immunological profiles *in vitro* (cytokines, chemokines, and growth factor secretion analyzed by flow cytometry) of a library of compounds derived from hot-water extracts of herbal medicines (Hioki et al., 2022).

**Figure 2** strikingly shows that the antibacterial and antileishmanial activities are correlated with two distinct groups of compounds. This is by no means surprising as these organisms have significantly different physiological organization, leading to different drugs currently used against these infections. The species showing the most significant antibacterial activities are *P. strigosum* and *P. xanthostachyum*, and the species showing the most significant antileishmanial activities are *P. pseudoarboreum*, *P. calvescentinerve*, *P. glabribaccum*, *P. reticulatum* and *P. sancti-felicis*. The compounds correlated with these activities within these extracts are rather specific to these species (**Figure 3**). An overall weak activity of these *Piper* extracts on *P. falciparum* (**Table 1**) is an unexpected result, as *Piper* species are usually reputed for their antimalarial potential.

Features from **Group 1** were active not only on bacteria but also on one fungus and one parasite. Although the high cross-activity observed for this group of compounds (**Figure 2** and **supp info 3**) could foretell of a lack of selectivity, these compounds did not cause cytotoxicity in mammalian cells. Six of the features of **Group 1** annotated as alkaloid derivatives were grouped in a same cluster in the molecular networking (MN), suggesting a similar biosynthesis pathway. Only three of them were annotated in our workflow, suggesting that the three unannotated ones may have an original structure. The reliability of the annotation or classification (NPClassifire and ClassyFire) part of the workflow may nevertheless be tempered, as shown by the case of feature 27, classified by the workflow as an alkaloid, but annotated as filfiline, a fatty amide previously isolated from the roots of *P. retrofractum* (Banerji et al., 2002). The broad bioactivity spectrum of alkaloids is well known, *i.e*., antibacterial, antiparasitic, anticancer, but also their cytotoxicity (Newman and Cragg, 2007, 2020; Daley and Cordell, 2021; Yan et al., 2021). The mechanism of action of antibacterial alkaloids is mainly described as disrupting the bacterial membrane, affecting DNA function and inhibiting protein synthesis (Ananthan et al., 2009; Pan et al., 2014; Kelley et al., 2013; Li et al., 2014; Larghi et al.). Antibacterial alkaloids (MIC <10 μM/mL) are as diverse as isoquinolines, aporphines, phenanthrenes, quinolines and indoles and their activity is mostly oriented against Gram-positive bacteria (Porras et al., 2021). Quinoleines like 8-hydroxyquinoline and its derivatives and evocarpine are active on bacteria associated with respiratory system infections: *M. tuberculosis*, *S. aureus*, and MRSA (Methicillin-resistant *Staphylococcus aureus*) (MIC ≤ 10 μM/mL). Aporphine alkaloid derivatives exhibit high broad-spectrum activity against Gram-positive bacteria: *S. agalactiae*, *S. aureus*, *S. epidermidis*, *E. faecalis*, and as in our study *E. coli* (Hamoud et al., 2015; Tan et al., 2015). Even if the chemical array within *Piper* spp. is impressively diverse, nitrogen-containing compounds are limited, *i.e*., alkaloids like piperines or amides like piplartine, phenethyls, and diaminodiamides. Amides are known to have fungicidal, cytotoxic, or antiprotozoal activities. With regard to fungicidal activity, dehydropipernonaline and nigramide R previously isolated from *P. retrofractum* displayed potent growth inhibition of *Cladosporium cladosporioides* and cytotoxicity against the L5178Y mouse lymphoma cell line (IC_50_ values of 8.9 μM and 9.3 μM, respectively) (Muharini et al., 2015). Amide piplartine isolated from a methanol extract of *P. retrofractum* showed good activity on *L. donovani* with an IC_50_ value at 7.5 μM (Bodiwala et al., 2007). In **Group 1**, features correlated with a fungicidal activity would only target *C. albicans*. Other active compounds identified in **Group 1** were one terpenoid, two fatty acids, one shikimate and phenylpropanoid derivative and one polyketide.

Compounds from the **Group 2**, characterized as being rather specific to *Leishmania*, are labeled as polyketides (3), shikimates and phenylpropanoids (3), terpenoids (2) and alkaloids (1). Their lack of activity on *Trypanosoma* is not surprising (**Figure 2**), as even though both parasites belong to the same *Trypanosomatidae* order, the current treatments are not similar. These two parasites cause different pathologies, in distinct geographical areas. They evolved differently and show evolutionary discrepancies in their mechanisms that may explain variations in sensitivity to treatments (Fernandes et al., 2020; Van den Broeck et al., 2020). A cross-reading of **Figures 2** and **3**, suggests that extracts of *P. calvescentinerve* and *P. glabribaccum* and features from **Group 2** have not only a selectivity for *Leishmania* but also a good selectivity index. Selectivity is a key parameter for antileishmanial drugs: antimonials, amphotericin B, paromomycin sulfate and miltefosine have variable efficacy against the 20+ *Leishmania* species but have significant adverse effects (Rao et al., 2019). Tremendous efforts have been put into the understanding of the *Leishmania* biology, leading to the identification of numerous putative targets: ergosterol and its biosynthetic pathway *[i.e*. amphotericine B and enzymes like squalene synthase (SQS) or sterol methyltransferase (SMT)], the glycolytic pathway necessary to provide glucose as an energy source, DNA topoisomerases, enzymes of the polyamine biosynthetic pathway (*i.e*. arginase, ornithine decarboxylase, s-adenosylmethionine decarboxylase, and spermidine synthase), redox metabolism pathway (Nagle et al., 2014; Raj et al., 2020).

The compounds labeled as shikimates or phenylpropanoids (acetanisole, aduncamide, and sesamin) or alkaloid (piperettine) were annotated at the genus level because they had previously been isolated from *P. tuberculatum*, *P. nigrum*, *P. longum*, *P. aduncum*, *P. puberulum*, *P. austrosinense*, *P. brachystachyum, P. mullesua, P. retrofractum, P. sarmentosum* and *P. sylvaticum* (Spring and Stark, 1950; Orjala et al., 1993; De Araujo-Junior et al., 1997; Puri et al., 1998; Umezawa, 2003; Tuntiwachwuttikul et al., 2006). Among these compounds, only the lignan sesamin was reported to show activity on *L. amazonensis* with an IC_50_ of 44.6 μM/mL and was not cytotoxic for mouse macrophage cells (CC_50_ > 100 μg/mL, SI > 6) (Pulivarthi et al., 2015). A study based on computational methods suggests that sesamin could be a promising inhibitor of the *L. donovani* CRK12 receptor (binding affinity of −8.5 kcal/mol) (Broni et al., 2021). Nevertheless, sesamin, isolated from a hexane extract of *P. retrofractum* (IC_50_ of the extract = 5 μg/mL), was inactive on *L. donovani* (Bodiwala et al., 2007) (Bodiwala et al., 2007). This compound was also inactive on *Plasmodium falciparum* K1 multidrug resistant strain*, Mycobacterium tuberculosis* H37Ra, and *Candida albicans* (EC50 > 20 μg/mL, MIC > 200 μg/mL and IC_50_ > 50 μg/mL, respectively). Cubein, another lignan isolated from *P. cubeba*, was shown to be active on *L. donovani* (IC_50_ = 28.0 μM). Interestingly, sesamin is connected in the MN with an unknown compound (RT = 6.577, [M+H]^+^ = 360.145). Piperettine has been previously isolated from *P. nigrum* and *P. aurantiacum* and was shown to be active on epimastigotes and amastigotes of *Trypanosoma cruzi* (IC_50_ = 10.67 and 7.40 μM, respectively) (Ribeiro et al., 2004). In our work, the compound annotated as piperettine was only detected in *P. calvescentinerve* extracts (**Figure 3**) but was not labeled as being correlated with any activity on *T. b. gambiense* (**supp info 3**). The use of this *Piper* species as a medicinal plant with various indications can partly be validated by the fact that it is an inhibitor of 5-lipoxygenase (76.02 μM) (Muthuraman et al., 2019), a key enzyme involved in the biosynthesis of pro-inflammatory leukotrienes, provided its concentration is sufficient in the traditional preparations. Aduncamide was shown to present a moderate antineuroinflammatory activity (IC_50_ at 26 ± 8.3 μM) by the Griess method on LPS-stimulated BV-2 cells (Zheng et al., 2021), to be cytotoxic for KB nasopharyngeal carcinoma cells (ED_50_ = 5.7 μg/mL) and to inhibit growth Gram-positive *B. subtilis* and *M. luteus*, while being less active towards Gram-negative *E. coli* (Orjala et al., 1993). In our work, the compound annotated as aduncamide was identified in extracts of *P. glabribaccum*, *P. calvescentinerve* and *P. cordatomentosa*, whose extracts are among the most active on *Leishmania*. Two compounds were labeled as 10-membered lactone macrolides and annotated as decarestrictin H and modiolide A, previously isolated from the fungi *Penicillium simplicissimum* and *Stagonospora cirsii*, respectively (Grabley et al., 1992; Evidente et al., 2008). These two last annotations, although resulting from similar fragmentation patterns, may be subject to caution. Indeed, these compounds were annotated at the generic level and no macrolide has been identified in *Piper* spp. so far (only a macrolactam, laevicarpin, previously isolated from *P. laevicarpus*) (da Silva A. Maciel et al., 2016). Furthermore, macrolides are well-known antibacterial compounds and thus should logically rather be clustered in **Group 1**, if ever present in the extracts. A compound annotated as apigenin was detected only in *P. xanthostachyum* and was identified as being correlated to the antibacterial activity (**Figure 2A**). This may appear surprising given that apigenin is a common and ubiquitous flavonoid with no antibacterial activity. The number of possible flavonoid isomers being quite large, this annotation may also be subject to caution, notwithstanding the fact that the actual compound (p87) being present in the extract remains of interest from the antibacterial perspective. Other compounds were annotated as flavonoids in the extracts (most of the compounds in **Group 3**, see **supp info 3**), characterized by a moderate activity of the extracts containing them, a moderate correlation with all the activities without any clear selectivity. These compounds were annotated at the genus level, as they were previously isolated from *Piper* species (Matsui and Munakata, 1976; Lago et al., 2004; González et al., 2022). Antiprotozoal or antimicrobial activity of flavonoids is well documented (Graf et al., 2005; Ortiz et al., 2017, 2020). Two compounds were annotated as pinocembrine and pinostrobin, both belonging to the flavanones subclass. Pinocembrine displays antifungal (*C. cladosporioides* and *C. sphaerospermum*) or antibacterial (*Enterococcus faecalis*, *Mycobacterium tuberculosis*) potential (Lago et al., 2004; Jeong et al., 2009; Gröblacher et al., 2012), while showing either no cytotoxicity on healthy and cancerous cell lines (RAW 264.7, epidermoid carcinoma of the oral cavity (KB), human small cell lung cancer (NCI-H187), metastatic murine colon 26-L5 carcinoma, PANC-1 human pancreatic cancer, metastatic human HT-1080 fibrosarcoma), or high cytotoxicity (Awale et al., 2009; Yenjai and Wanich, 2010; Lee et al., 2013). Pinostrobin showed weak antimalaria activity on *P. falciparum* (Kaur et al., 2009) but significant potential as a cancer chemopreventive agent (Gu et al., 2002).

Some of these *Piper* compounds are therefore of interest in an isolation and factual testing perspective. Compounds annotated as aduncamide, sesamin, and apigenin, along with the unknown compounds correlated to the activity, could be subjected to mass-targeted isolation (Vásquez-Ocmín et al., 2022a), in order to confirm their annotation or identify their structure, as well as confirm their biological potential.

## CONCLUSION

Hyphenated analytical techniques are very useful to provide highly informative chemical profiling of complex metabolomes. Nevertheless, deciphering such profiles to determine the compounds responsible for the biological activity remains a challenge. By correlating these chemical profiles with biological assay results, we propose a workflow of integrated metabolomics statistical tools to provide a broad picture of the metabolome as well as a map of compounds putatively associated to the bioactivity. The structure of these compounds can either be annotated from previous works or remain unknown at this stage, but in any case, they require a formal isolation step to confirm their structure and activity. Nevertheless, an initial data mining on analytical-scale data proves very effective for prioritizing the compounds to target. The relevance of our approach has been validated on a set of *Piper* species tested for their anti-infectious diseases at the extract level. Such an approach can be extended to any type of natural extract, particularly when prior phytochemical data is available in the literature.

## Supporting information

Deciphering anti-infectious compounds from Peruvian medicinal Cordoncillos-SI_Vasquez-Ocmin

## DATA AVAILABILITY STATEMENT

The raw data from the LCMS were uploaded to zenodo (DOI:10.5281/zenodo.7317966).

## AUTHOR CONTRIBUTIONS

Conceptualization: PGVO, AM, GM, and LRM; Methodology: PGVO, AM, GM, and LRM; Chemical data acquisition/curation: PGVO, KL, GM, AG, and AM; Statistical analysis: PGVO and GM; Biological data curation: SC, VR, SB, and SP; Investigation: PGVO, SC, VR, GM, SP, AG, KL, ID, LRV, HRC, WRM, KL, SB, LRM, and AM. PGVO wrote the original draft and all the authors contributed to the final manuscript.

## ACKNOWLEDGMENTS

Authors are indebted to the communities cited in this work for their essential contribution in order to spread their ancestral knowledge. The authors thank IRD the French National Research Institute for Sustainable Development (IRD), UMR 152 PHARMADEV, IRD-UPS (Toulouse, France) for the post-doctoral fellowship to PG V-O (contract number 04077858). The authors thank MINEDU (Ministerio de Educación), Lima, Peru, convenio MINEDU-UNAP (Decreto Supremo 146-2020, EF/Resolución Rectoral N°1312-2020-UNAP, 9/12/2020). We also appreciate the important collaboration of Juan Celedonio Ruiz (Herbarium Amazonense (AMAZ) de la Universidad Nacional de la Amazonía Peruana (UNAP), Iquitos, Perú). The authors are also grateful to Amélie Perez (MetaboHUB-MetaToul-Agromix, LRSV, Auzeville-Tolosane), Guillaume Cabanac (IRIT, Toulouse), and Rizwana Zaffaroullah (CNR du Paludisme, AP-HP, Hôpital Bichat - Claude Bernard, F-75018 Paris, France).

