## Supplementary material for "Deciphering anti-infectious compounds from Peruvian medicinal *Cordoncillos* extract library through multiplexed assays and chemical profiling": Deciphering anti-infectious compounds from Peruvian medicinal Cordoncillos-SI_Vasquez-Ocmin

[†]Curent affiliation: Laboratoire de Recherche en Sciences Végétales (UMR 5546), CNRS, Université de Toulouse, Toulouse, France.
[‡]These authors contributed equally to this work
[⊥]These authors share senior authorship
[1]UMR152 PHARMADEV, IRD, UPS, Université de Toulouse, Toulouse, France
[2]Université Paris-Saclay, CNRS, BioCIS, 91400, Orsay, France
[3]CNR du Paludisme, AP-HP, Hôpital Bichat - Claude Bernard, F-75018 Paris, France
[4]Joint Research Unit 1158 BioEcoAgro, Univ. Lille, JUNIA, INRAE, Univ. Liège, UPJV, Univ. Artois, ULCO, F-59650, Villeneuve d'Ascq, France
[5]Laboratoire de Recherche en Sciences Végétales (UMR 5546), CNRS, Université de Toulouse, Toulouse, France
[6]MetaboHUB, National Infrastructure of Metabolomics and Fluxomics, Toulouse, France
[7]Facultad de Farmacia y Bioquímica, Universidad Nacional de la Amazonía Peruana (UNAP), Iquitos 16000, Peru
[8]Centro de Investigación de Recursos Naturales, Universidad Nacional de la Amazonía Peruana (UNAP), Iquitos 16002, Peru
[9]Facultad de Ingeniería Química, Universidad Nacional de la Amazonía Peruana (UNAP), Iquitos 16002, Peru
[10]International Joint Laboratory of Molecular Anthropological Oncology (LOAM), National Cancer Institute, Lima, Perú

*** Correspondence:**
Pedro G. Vásquez-Ocmín

Alexandre Maciuk


### Abstract

High prevalence of parasitic or bacterial infectious diseases in some world areas is due to multiple reasons, including a lack of an appropriate health policy, challenging logistics and poverty. The support to research and development of new medicines to fight infectious diseases is one of the sustainable development goals promoted by World Health Organization (WHO). In this sense, the traditional medicinal knowledge substantiated by ethnopharmacology is a valuable starting point for drug discovery. This work aims at the scientific validation of the traditional use of *Piper* species as firsthand anti-infectious medicines. For this purpose, we adapted a computational statistical model to correlate the LCMS chemical profiles of 19 *Piper* species to their corresponding anti-infectious assay results based on 37 microbial or parasites strains. We mainly identified two groups of bioactive compounds (called features as they are considered at the analytical level and are not formally isolated). Group 1 is composed of 11 features being highly correlated to an inhibiting activity on 21 bacteria (principally Gram-positive strains), one fungus (*C. albicans*), and one parasite (*Trypanosoma brucei gambiense*). The group 2 is composed of 9 features having a clear selectivity on *Leishmania* (all strains, both axenic and intramacrophagic). Bioactive features in group 1 were identified principally in the extracts of *Piper strigosum* and *P. xanthostachyum*. In group 2, bioactive features were distributed in the extracts of 14 *Piper* species. This multiplexed approach provided a broad picture of the metabolome as well as a map of compounds putatively associated to bioactivity. To our knowledge, the implementation of this type of metabolomics tools aimed at identifying bioactive compounds has not been used so far.

**Keywords:** Anti-infectious diseases, *cordoncillos*, metabolomics, Peruvian Amazonia, *Piper*

**Supporting information 1.** Set samples and description of *Piper* species

| Date of collection | Species | Part extracted | Voucher number | UTM coordinates | Place of collection, department |
|---|---|---|---|---|---|
| 14/12/2020 | *Piper casapiense* (Miq.) C. DC. | Leaves and stems | 041044 | 755672 8111253 | Estación Isula-Rio Napo, Lago Sunimiraño, Loreto |
| 14/12/2020 | *Piper strigosum* Trel. | Leaves and stems | 029675 | 513164,15 8111231 | Estación Isula-Rio Napo, Loreto |
| 01/08/2020 | *Piper pseudoarboreum* Yunck | Leaves and stems | 019724 | 526944 8141995 | Carretera Iquitos Nauta km 29, distrito de San Juan Bautista, Loreto |
| 27/11/2020 | *Piper armatum* Trel. & Yunck | Leaves and stems | 024563 | 527171 8118136 | Comunidad del gallito, Rio Amazonas, Loreto |
| 11/12/2010 | *Piper brasiliense* C. DC | Leaves and stems | 033310 | 0711335 9623600 | Estación Isula-Rio Napo, Lago Sunimiraño, Loreto |
| 6/12/2020 | “*Piper bullatum* Vahl” | Leaves and stems | 23129 | 295957 8557659 | Ámbito del valle de Kosñipata, distrito de Kosñipata, Provincia San Pedro, Cusco |
| 6/12/2020 | *Piper calvescentinerve* Trel | Leaves and stems | 23127 | 233321 8559681 | Ámbito del valle de Kosñipata, distrito de Kosñipata, Provincia San Pedro, Chontachaca, Cusco |

| | | | | | |
|---|---|---|---|---|---|
| 6/12/2020 | *“Piper cordatomentosa"* | Leaves and stems | 23125 | 233321<br>8559681 | Ámbito del valle de Kosñipata, distrito de Kosñipata, Provincia San Pedro, Chontachaca, Cusco |
| 6/12/2020 | *Piper crassinervium* Kunth. | Leaves and stems | 23128 | 295957<br>8557659 | Ámbito del valle de Kosñipata, distrito de Kosñipata, Provincia San Pedro, Chontachaca, Cusco |
| 22/09/2010 | *Piper divaricatum* G. Mey | Leaves | 10538 | 0288151<br>9312640 | Comunidad nuevo Cutervo, distrito Jepelacio, San Martin |
| 6/12/2020 | *Piper glabribaccum* Trel | Leaves and stems | 23121 | 215490<br>8544039 | Ámbito del valle de Kosñipata, San Pedro, Cusco |
| 20/12/2009 | *Piper heterophyllum* Ruiz & Pav. | Leaves | 028164 | 513560<br>8111901 | Carretera Mazan-Indiana, Loreto. |
| 6/12/2020 | *Piper oblongum* Kunth | Leaves and stems | 23123 | 231733<br>8559990 | Ámbito del valle de Kosñipata, distrito de Kosñipata, Provincia San Pedro, Buenos aires, Cuzco. |
| 11/01/2010 | *Piper reticulatum* L | Leaves | 042127 | 531366<br>8146976 | Carretera Iquitos – Nauta, Provincia Maynas , Loreto |
| 07/11/2012 | *Piper sancti-felicis* Trel | Leaves and stems | 006367 | 0713020<br>9622125 | Estación Isula-Rio Napo, Lago Sunimiraño, Loreto |

| | | | | | |
|---|---|---|---|---|---|
| 14/12/2020 | *Piper stellipilum* (Miq.) C. DC | Leaves and stems | 039893 | 513250<br>8111253 | Estación Isula-Rio Napo, Lago Sunimiraño, Loreto |
| 6/12/2020 | *Piper trigonum* C. DC. | Leaves and stems | 23124 | 231733<br>8559990 | Ambito del valle de Kosñipata, distrito de Kosñipata, Provincia San Pedro, Chontachaca, Cusco |
| 6/12/2020 | *Piper verruculosum* C. DC. | Leaves and stems | 23122 | 215490<br>8544039 | Ámbito del valle de Kosñipata, Buenos Aires, Cusco |
| 29/12/2009 | *Piper xanthostachyum* C.DC. | Leaves | 10491 | 526941<br>8141988 | Alpahuayo Mishana, Carretera Iquitos - Nauta, Provincia Maynas, Loreto. |

**Supporting information 3.** Clustered Image Map (CIM) performed on the two-blocks of data sets (*in extenso*)

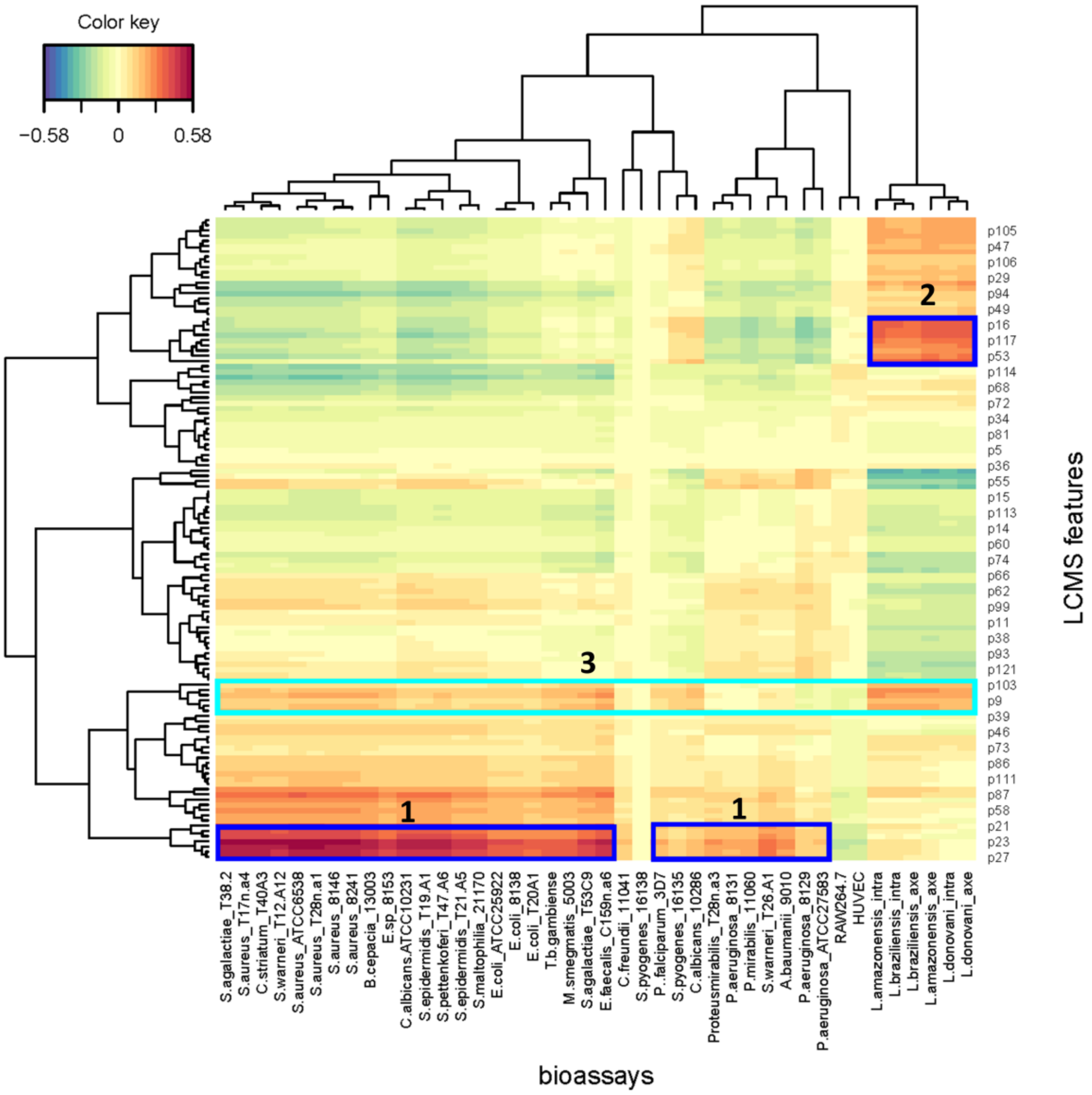
